# Causal effects of prefrontal transcranial magnetic stimulation on dopamine-mediated reinforcement learning in healthy adults

**DOI:** 10.1101/2022.06.03.494692

**Authors:** Kathryn Biernacki, Catherine E. Myers, Sally Cole, James F. Cavanagh, Travis E. Baker

**Author notes:** **Corresponding author**: Kathryn Biernacki PhD, Center for Molecular and Behavioral Neuroscience, Rutgers University – Newark, 197 University Ave, Newark NJ, 07302. **Declarations of interest:** none.

## Abstract

**Background:** 10-Hz repetitive transcranial magnetic stimulation (rTMS) to the left dorsal lateral prefrontal cortex (DLPFC) has been shown to increase dopaminergic activity in the dorsal striatum, a region strongly implicated in reinforcement learning. However, the behavioural influence of this effect remains largely unknown.

**Objective:** Here, we tested the causal effects of rTMS on behavioral and computational characteristics of reinforcement learning.

**Methods:** 40 healthy individuals were randomized into Active and Sham rTMS groups. Each participant underwent one 10-Hz rTMS session (1500 pulses) in which stimulation was applied over the left DLPFC using a robotic arm. Participants then completed a reinforcement learning task sensitive to striatal dopamine functioning. Participants’ trial-to-trial training choices were modelled using a reinforcement learning model (Q-learning) that calculates separate learning rates associated with positive and negative reward prediction errors.

**Results:** Subjects receiving Active TMS exhibited an increased reward rate (number of correct responses per second of task activity) compared to the Sham rTMS group. Computationally, the Active rTMS group displayed a higher learning rate for correct trials (αG) compared to incorrect trials (αL). Finally, when tested with novel pairs of stimuli, the Active group displayed extremely fast reaction times, and a trend towards a higher reward rate.

**Conclusions:** The present study provided specific behavioral and computational accounts of altered striatal-mediated reinforcement learning induced by a proposed increase of dopamine activity by 10-Hz rTMS to the left DLPFC. Together, these findings bolster the use of TMS to target neurocognitive disturbances attributed to the dysregulation of dopaminergic-striatal circuits.

## Introduction

Positron emission tomography (PET) studies have demonstrated that 10-Hz repetitive transcranial magnetic stimulation (TMS) to the left dorsal lateral prefrontal cortex (DLPFC) can modulate dopaminergic activity in the dorsal striatum (caudate, but not putamen) and prefrontal cortex of healthy subjects [1–3]. Current thinking holds that stimulation of the left DLPFC subsequently stimulates midbrain dopamine neurons, which in turn enhances dopamine release in its projected targets (e.g., striatum, prefrontal cortex)[2], an argument supported by a recent TMS and optogenetic study[4–6]. It has been suggested that the observed modulation of dopamine release might mediate, at least in part, the therapeutic effects of TMS seen in clinical populations, such as patients with depression and substance use disorders[7–13]. Although current evidence details the enhancing effects of TMS on striatal-dopamine activity, it remains unclear whether TMS has a causal effect on the putative function of the dorsal striatum.

Phasic midbrain dopamine activity encodes a reward prediction error (RPE) signal [14, 15], which guides stimulus and action value learning as assessed by reinforcement learning models [16, 17]. Accordingly, an influential neurocomputational model of reinforcement learning, the “basal ganglia go/no-go model,” holds that phasic bursts of dopamine activity (positive RPEs) following unexpected positive feedback facilitate approach or ‘go’ learning by reinforcing dorsal striatal D1 receptors, whereas transient cessations in dopamine activity (negative RPEs) following unexpected negative feedback facilitate avoidance or ‘no-go’ learning by reinforcing dorsal striatal D2 receptors [18]. Extensive psychiatric and neurological research indicate that a disruption in positive and negative RPE signaling in the dorsal striatum can selectively impair approach and avoidance learning, respectively [18, 19]. For example, the model predicts that people with Parkinson’s disease will be more accurate at approach than avoidance learning while on dopaminergic medication due to an enhanced phasic dopamine burst, and the reverse while off medication due to a diminished phasic dopamine pause, findings that have been confirmed empirically [18, 20]. These studies suggest that a TMS-induced enhancement of dopamine signaling in the dorsal striatum should be paralleled by alterations in approach and avoidance learning, as well as computational variables associated with reinforcement learning[21].

To test this hypothesis^a^, we examined the impact of 10-Hz rTMS to the left DLPFC on behavioral and computational characteristics of the striatal dopamine system described by reinforcement learning theory. Here, we randomly assigned subjects into an Active and Sham TMS group, and then following 10-Hz rTMS to the left DLPFC, participants engaged in the Probabilistic Selection Task (PST), a trial-and-error learning task shown to be sensitive to disturbances in striatal dopamine signaling [18]. Accordingly, we predicted that if 10-Hz rTMS differentially affects dopaminergic RPE signals during reinforcement learning, then performance in this task should differ between Active and Sham TMS groups. Specifically, we predicted that subjects receiving Active TMS will be more accurate at approach learning due to an enhanced dopamine signal. Furthermore, to examine computational differences in reinforcement learning across rTMS groups, we simulated PST performance using a reinforcement learning model known as Q-learning, which provides a means to simulate individual trial-to-trial task choices to determine separate learning rates for positive and negative RPE signals [17, 27, 28]. The gain (αG) and loss (αL) learning rates determine the degree to which recent RPEs affect the expected value [17]. Thus, if RPE signals are artificially amplified in response to TMS, then changes in gain and loss learning rates should capture such TMS-induced RPE alterations.

## Methods

### Participant recruitment and study, procedures

A total of 44 healthy participants were recruited and assessed for contraindications to TMS (e.g., epilepsy, metal implants), current use of central nervous system drugs, neurological disorders, and psychiatric symptoms (e.g., depression, anxiety, substance use disorders). All participants had normal or corrected-to-normal vision and were naive about the purpose of the study. Of these 44 participants, 5 participants met exclusion criteria due to TMS contraindications. Four participants who failed to learn the PST task were excluded from the TMS analysis (see Probabilistic Selection Task section). In total, the data of 35 healthy controls were included in this study, with 18 subjects for the TMS Active group and 17 subjects for the Sham group (see Table 1 for subject/group information). The study was approved by the local research ethics committee at Rutgers University, USA, and was conducted in accordance with the ethical standards prescribed in the 1964 Declaration of Helsinki. Participants were compensated at $75 USD for completing the three-hour study.

**Table 1.**
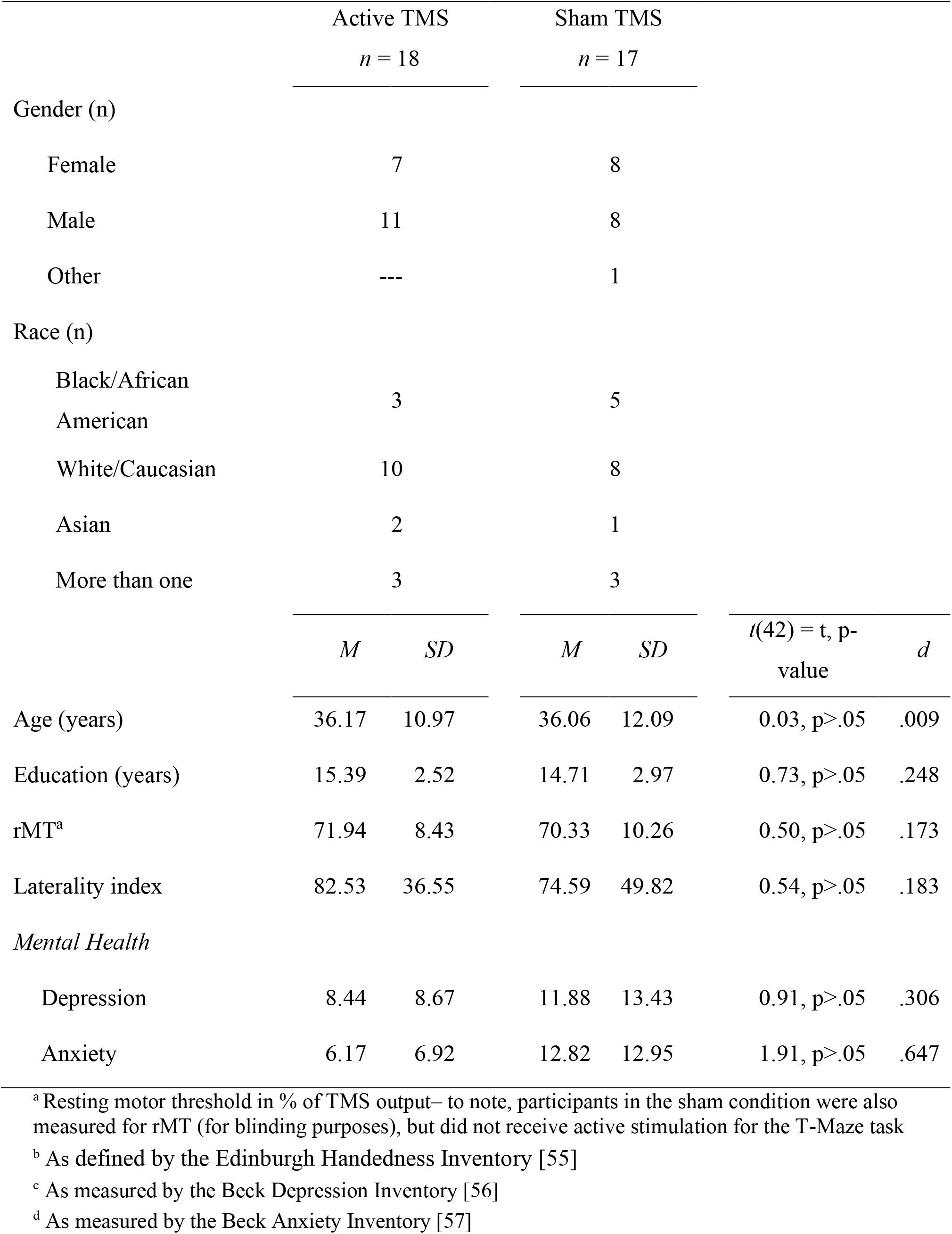
Participant Characteristics

### Probabilistic selection task (PST)

Participants completed an adapted version^b^ of the PST (Figure 1B). The PST consisted of a forced choice training phase consisting of three blocks of sixty trials each, followed by a subsequent testing phase [18]. During the training phase, the participants were presented with two stimulus pairs, where each stimulus was associated with a different probabilistic chance of receiving “Correct” or “Incorrect” feedback. These stimulus pairs (and their reward probabilities) were termed A/B (80%/20%) and C/D (70%/30%). Over the course of the training phase, a participant usually learns to choose A over B, and C over D, solely due to adaptive responding based on the feedback. During the testing phase, participants were exposed to all possible combinations of these stimuli (i.e., AB, CD, AC, AD, BC, BD) in a random order and were required to select the symbol in each pair that they believed to be correct, but without receiving any feedback about their choices. Each test pair was presented ten times. Data from the test phase were only analysed if participants selected the most rewarding stimulus (A) over the least rewarding stimulus (B) more than chance (≥60%) of the time when this stimulus pair was presented during the testing phase, since data from participants who fail this basic criterion are not interpretable [28–30]. This criterion removed 4 participants from the analyses, consistent with previous work [30]. For the remaining participants, accuracy for “approach learning” was scored as selecting the most-often rewarded A stimulus when paired with stimuli C or D, and for “avoidance learning” as not selecting (avoiding) the least-often rewarded B stimulus when paired with C or D.

**Figure 1.**
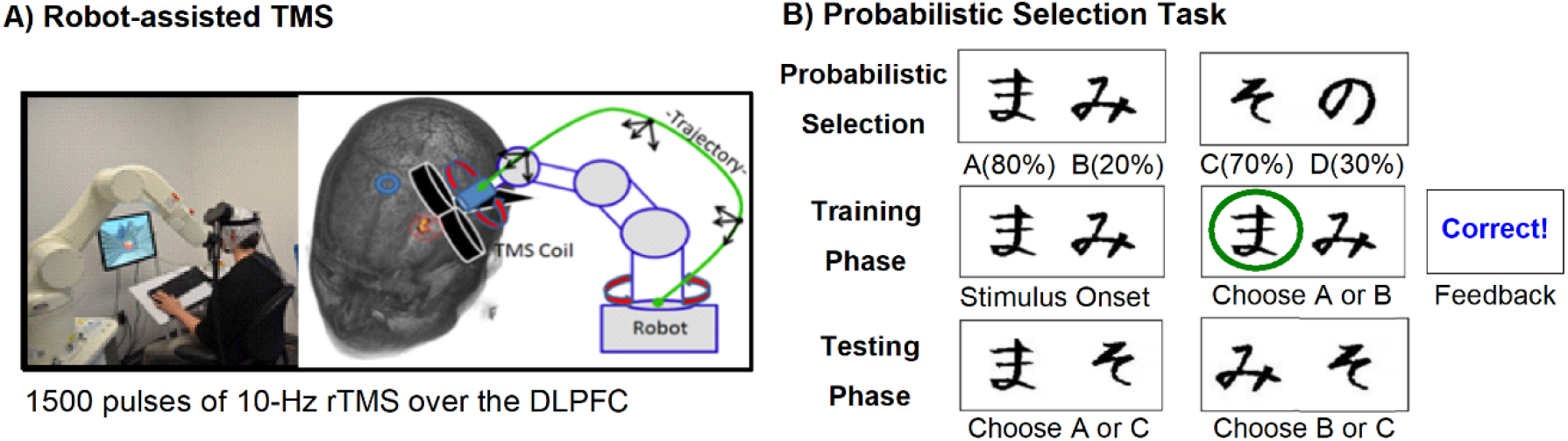
Experiment set-up. A) Robot-assisted repetitive transcranial magnetic stimulation (rTMS), to position the TMS coil directly over the participant’s dorsolateral prefrontal cortex (DLPFC). B) Top: Stimuli and outcome categories in the Probabilistic Selection Task. Middle, Bottom: Schematic of example trials during the training and testing phase.

### Q-learning Model

The trial-by-trial sequence of choices on the PST for each subject was fit using a Q-learning reinforcement learning model [17, 27], as detailed in [28]. Briefly, this model assigns expected reward values to actions taken during a particular state (i.e., choosing action A when seeing an A/B stimulus pair). These state-action values are referred to as Q values. The model used separate learning rate parameters for gain (correct, α_G_) and loss (incorrect, α_L_) feedback trials in the training phase of the PST, and these separate learning rates scaled the updating of the Q values separately for rewards and punishments. The expected value (*Q*) of any stimulus (*i*) at time (*t*) was updated after each reinforcer (*R*(*t*)=1 for correct, *R*=0 for incorrect):

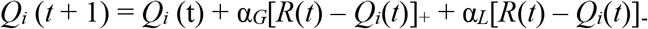

where the learning rate applied is either α_*G*_ or α_*L*_ depending on whether the outcome is better or worse than expected, respectively. These Q values were entered into a softmax logistic function to produce probabilities (*P*) of responses for each trial, with higher probabilities predicted for stimuli having relatively larger Q values, and a free parameter for inverse gain (*β*) to reflect the tendency to explore or exploit:

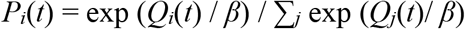

These probabilities were then used to compute the log likelihood estimate (LLE) of the subject having chosen that set of responses for a given set of parameters over the whole training phase. The parameters that produce the maximum LLE were found using the Simplex method, a standard hill-climbing search algorithm implemented with Matlab (The MathWorks, Natick, MA) function “fmincon”, thus providing a model fit for each individual subject that maximized the likelihood of producing the actions selected by the subject:

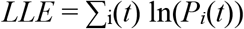

The parameters that produced the maximum log likelihood were selected and the Q values produced by this particular set of parameters were saved. Characterization of model fits were computed as pseudo-R^2^ statistics ((LLE-*r*)*/r*, where *r* is the LLE of a chance performance model). For further details on Q-learning and model fits, see [28].

### Robot-guided transcranial magnetic stimulation

rTMS was applied according to safety guidelines based on available safety studies of rTMS [31], and using a MagPro X100 stimulator with the Cool-B70 figure-of-eight coil (MagVenture, Falun, Denmark). The resting motor threshold (rMT) was determined individually before the Active and Sham stimulations. The stimulation intensity was 110% of the rMT of the right abductor pollicis brevis muscle. Average rMT stimulation intensity was 71.21% (range 50%-80%) and maximum stimulation output was 88% (range: 55%–88%). At the start of the experiment a 3D model of the subject’s head surface was created, and the robotic arm, head model, and tracking system were registered to a common coordinate system (SmartMove, ANT Neuro). The TMS target was placed on the x y z coordinate corresponding to the F3 position of the 10–20 EEG reference system, and the corresponding point on the surface was selected and the coil orientation was set at 45° angle. The virtual coil position was then transformed to robot coordinates, which define the movement of the robot to the corresponding target position relative to the subject’s head. The position of the cranium is continuously tracked, and the trajectory of the coil’s path adapted to head movements in real-time. This ensures a precise <1 mm coil-to-target position and allows free head movements during the experiment[32].

Prior to the PST task, the robotic arm positioned the TMS coil over the left DLPFC target, electrode F3. Each participant received a single session of stimulation which consisted of 1,500 pulses of rTMS (10 Hz trains for 5 sec; repeated 30 times with an inter-train interval of 30 sec; total session, 30 minutes). The Active and Sham rTMS procedures were performed at the same location on the skull; however, for the Sham stimulation, the coil was flipped 180° to ensure that participants did not receive active simulation but received the same auditory sensation. To note, during the 30 sec inter-train TMS interval, participants completed 10 trials of a virtual T-maze task, and EEG was recorded throughout the T-maze task and PST as part of a TMS study on reward processing and opioid use disorder, the results of which will be reported elsewhere. After the TMS session was completed, participants engaged in the PST task within 5min of the last pulse sequence. At the end of the experiment, participants were debriefed and compensated for their participation.

## Results

### Training phase

We first analysed overall PST training phase accuracy and reaction time using a repeated measures ANOVA with Block (BLK1, BLK2, BLK3) and Stimulus (AB, CD) as within-subject factors, and TMS group (Active vs Sham) as a between-group factor (Figure 2A and 2B, left panel). Regarding accuracy, a main effect of Stimulus was observed, *F*_1, 33_ = 4.49, *p* <.05, η_p_^2^ =0.13, indicating that subjects performed better on AB trials (*M* = 76%, *SEM* = 2) compared to CD trials (*M* = 69%, *SEM* = 2) (Figure 2B). No main effect of Block or TMS group, nor an interaction, was observed (*p*>.05). For descriptive purposes, we included the results of the stimulus specific training phase data (Figure 2B). An exploratory within-group analysis revealed a main effect of Block for the Active TMS group, *F*_2, 34_ = 4.7, p <.01, η_p_^2^ =0.22, indicating that relative to Block1 (*M* = 71%, *SEM* = 3), participants selected the most optimal stimulus more often at Block 2 (*M* = 77%, *SEM* = 4, p < .05) and Block 3 (*M* = 78%, *SEM* = 3, p < .01) (Figure 2A). Regarding reaction time (Figure 2A and 2B, right panel), a main effect of TMS group was observed, *F*_1, 33_ = 7.92, *p* <.01, η_p_^2^ =0.20, indicating that the Active TMS group responded faster (*M* = 671 ms, *SEM* = 80) compared to the Sham TMS group (*M* = 996ms, *SEM* = 82, Figure 2A, right). This analysis also revealed an interaction between Stimulus and Block, *F*_2, 66_ = 4.1, *p* <.05, η_p_^2^ =0.11, indicating that reaction time increased across blocks for the CD pair, but not the AB pair.

**Figure 2.**
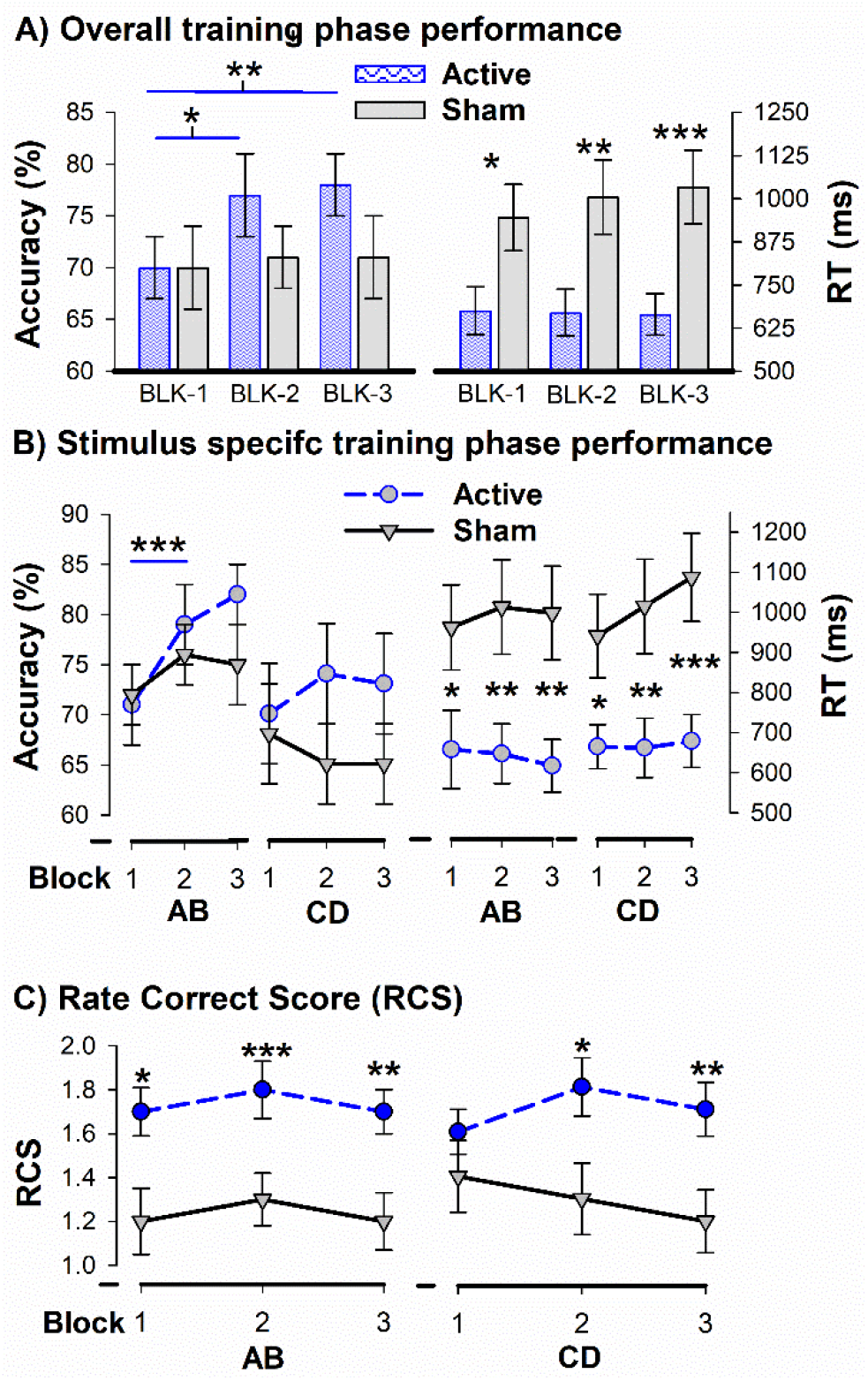
Training phase performance. **A)** Overall and **B)** stimulus specific accuracy (% correct) and reaction time, RT (ms), across blocks of the PST for the active (blue) and sham (grey) rTMS groups. **C)** Rate Correct Score (RCS; number of correct responses produced per second of task activity), indicating active (blue) and sham (grey) group. * = *p* < .05, ** = *p* < .01, *** = *p* < .005. Error bars indicate standard errors of the means.

Next, in order to take speed as well as accuracy into account, we calculated a combined measure called the Rate Correct Score [RCS; 33]. The RCS is the number of correct responses divided by the sum of correct and incorrect response times and is easily interpretable as the number of correct responses produced per second of task activity (i.e., reward rate) [33]. RCS recovers the information from accuracy and response times and can account for a larger proportion of the variance than the separate measures. In a comparison of several methods that combined accuracy and response times, RCS was found to be preferable to other measures [34]. This analysis revealed a main effect of Group, *F*_1, 33_ = 6.67, *p* <.01, η_p_^2^ =0.17, indicating that the Active rTMS group exhibited a higher RCS score (RCS, *M* = 1.7 correct responses per second, *SEM* = .11) relative to the Sham rTMS group, (RCS, *M* = 1.3 correct responses per second, *SEM* = .11). In other words, participants receiving active rTMS showed a greater reward rate compared to the Sham group. A 3-way interaction was also observed, indicating that for the CD trials, the reward rate increased for the Active TMS group across blocks, but decreased for the Sham TMS group across blocks (Figure 2B).

### Testing Phase

A repeated-measures ANOVA on test phase accuracy with stimulus condition (approach and avoidance) as a within-subject factor and TMS group (Sham vs. Active) as a between-subject factor did not reveal a main effect nor an interaction, (*p*>.05) (Figure 3A, left). To note, an exploratory analysis on stimulus specific conditions revealed a trend, such that active TMS participants displayed higher accuracy rates for AD pairs compared to Sham, p = .06 (Figure 3B). In regard to RT, a main effect of condition was observed, *F*_1, 33_ = 10.97, *p* < .005, η_p_^2^ =0.25, indicating that subjects performed faster at approaching A (*M* = 962ms, *SEM* = 61) compared to avoiding B (*M* = 1136ms, *SEM* = 87) (Figure 3A, right). Further, a main effect of group was also observed, *F*_1, 33_ = 4.55, *p* < .05, η_p_^2^ =0.12, indicating that the Active TMS group performed faster (*M* = 898ms, *SEM* = 98) compared to the Sham group (*M* = 1200ms, *SEM* = 101). A subsequent analysis indicated that this main effect of group was largely driven by the approach condition (*p*<.01) and not by the avoidance condition (*p* = .09) (Figure 3A, left panel). In regards to the RCS analysis, no main effects nor interaction were observed (all *p*’s>.05) (Figure 3C). An exploratory analysis revealed the Active TMS group exhibited a higher RCS score for the approach condition (RCS, *M* = 1.4, *SEM* = .11) relative to avoidance condition (RCS, *M* = 1.2, *SEM* = .11), *t*_(17)_2.4, *p* < .05. Further, the results revealed a trend for a group difference in reward rate for the approach condition (Active rTMS: RCS, *M* = 1.4, *SEM* = .11 | Sham rTMS: RCS, *M* = 1.1, *SEM* = .11), *t*_(33)_1.9, *p* = .07.

**Figure 3.**
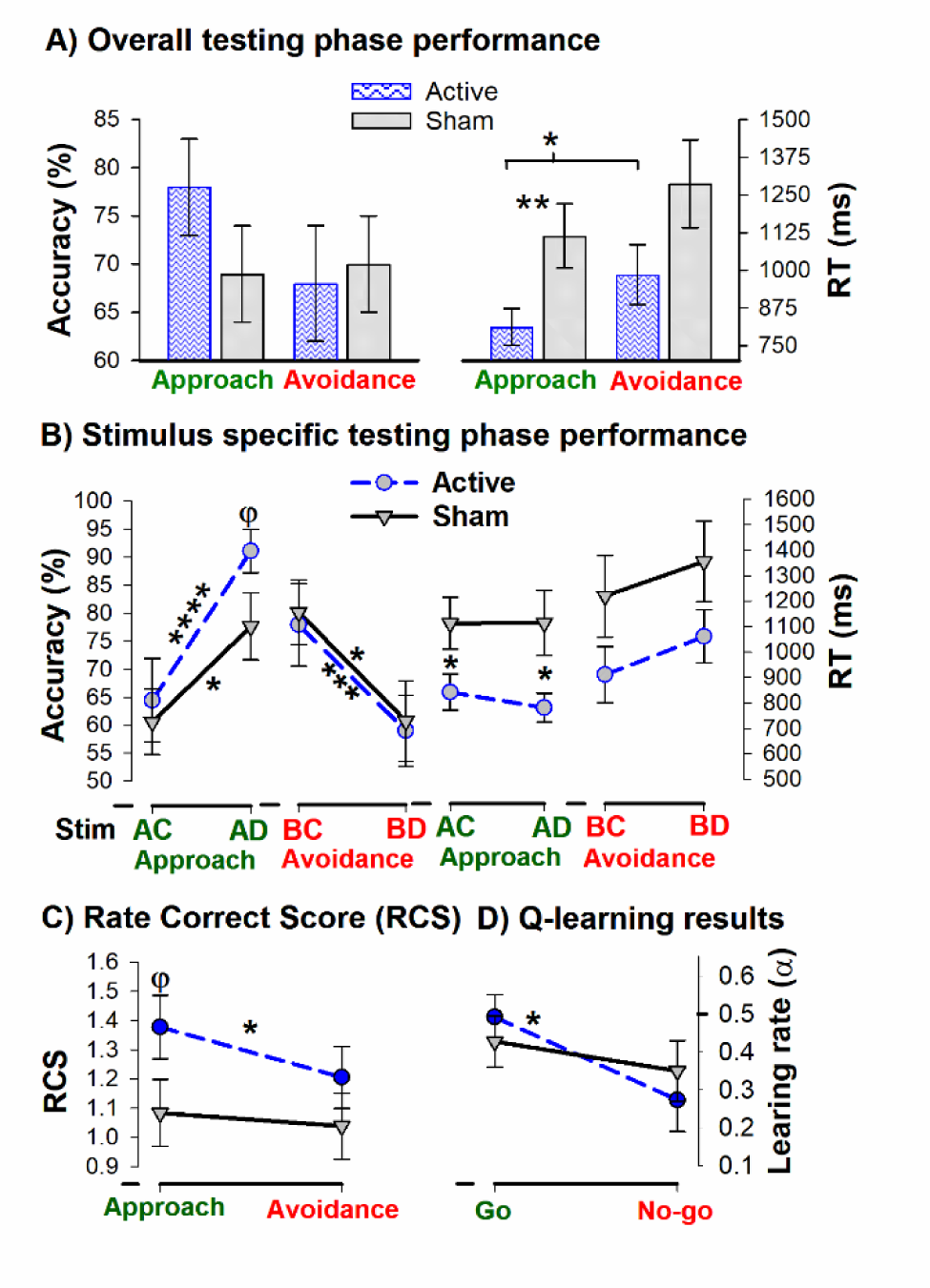
Test phase and computational results. **(A)** Overall and **(B)** stimulus specific test phase accuracy (left panel) and RT (Right panel) data for the Active (blue) and Sham (grey) groups, separately for the Approach and Avoidance conditions. **C)** Rate Correct Score (RCS) for Approach and Avoidance conditions, for active (blue dashed lines) and Sham (black solid lines) rTMS groups. **D)** Q-learning results. Go and No-Go learning rate values for Active (blue dashed lines) and Sham (black solid lines) rTMS groups. Asterisk denotes significance of paired (within-group) and independent (between-group) t-test results. * = *p* < .05, ** = *p* < .01, *** = *p* < .005, **** p < .001. φ = trend, p<.07. Error bars indicate standard errors of the mean.

### Q-learning Results

Average log likelihood for the training fit was 96 (*SEM*= 4.3), and average pseudo-R^2^ was .23 (*SEM* = .04). A repeated-measures ANOVA on average learning rate for correct trials (α_Gain_) and incorrect trials (α_Loss_) did not reveal a main effect nor an interaction (all *p’s* >.05). Next, we examined whether the modelling data were in line with the behavioral effects observed within the active rTMS group. Indeed, this within-group analysis revealed a shift towards higher gain learning rates (α_G_) relative to loss learning rates (α_L_) in the active rTMS group, *t*_(33)_=2.1, *p*<.05, an observation not observed in the Sham rTMS Group (Figure 3D). No group differences were observed for the inverse gain (β) parameter (*p*>.05, average value = 3.2, SEM = .34).

## Discussion

Dopamine RPE signaling in the dorsal striatum —observed both in the firing patterns of dopamine neurons and in dopamine concentration changes in the striatum —is essential for reinforcement learning [34–36]. While this proposal has been directly tested in animals using optogenetics [35], pharmacological manipulations of dopaminergic RPE signaling in the dorsal striatum rigorously in humans has been difficult (e.g., dopamine drugs have differing effects on various brain regions and reward phases [37], and no D1 agonist drugs are available for human use). Motivated by evidence demonstrating that 10-Hz rTMS to the left DLPFC can enhance dopaminergic activity in the dorsal striatum [3], we investigated the impact of rTMS on behavioral and computational measures of striatal-based reinforcement learning in healthy subjects.

Foremost, participants receiving active rTMS displayed a higher reward rate of task activity (RCS) during the training phase compared to those receiving Sham rTMS. Further, training phase accuracy (percent choice of the optimal action) increased across task blocks for the active rTMS group, but not the Sham rTMS group, suggestive of an increase in precision. Considerable evidence has shown that phasic bursts in dopamine activity encode positive RPE signals [16, 18, 34, 38], and it has been proposed that positive RPEs facilitate reward learning in the PST by reinforcing dorsal striatal connections [18, 38–40]. In parallel, 10-Hz rTMS to the left DLPFC has been shown to enhance dopamine release in the striatum [3]. By extension, we hypothesized that rTMS would be expected to increase positive RPE signals in the striatum and thereby enhance reward learning. Consistent with this hypothesis, it is reasonable to assume that the enhancement in learning accuracy across blocks and higher reward rate of task activity (RCS) observed in the Active rTMS group was mediated by a TMS-induced potentiation of dopamine RPE signaling in the dorsal striatum. Moreover, in the Q-learning model, action values are updated by the product of RPE and the learning rate. Thus, our observation that subjects receiving active rTMS displayed a higher gain learning rate relative to loss is consistent with the predictions of the go/no-go model. Specifically, by increasing phasic responsiveness of dopamine neurons, rTMS could act to increase the gain on the positive RPE signal in the striatum, and subsequently enhance positive learning rates in the model. Consistent with this interpretation, low-dose D2 receptor antagonism (presumed to increase striatal dopamine) [41, 42], and cigarette consumption [43] have also been shown to enhance positive RPE signaling, and subsequently reward learning in the PST.

While the RPE hypothesis of dopamine may provide one account of these findings, alternative explanations exist. Foremost, like Parkinson’s patients on dopamine medication, we predicted that subjects receiving active TMS will be more accurate at approach learning in the PST test phase due to an enhanced dopamine signal during learning. In contrast to this prediction, no group differences were observed on PST test phase performance and model-derived learning rates. Instead, the most robust group effect was observed in response behavior, such that 10-Hz rTMS hastened reaction time without compromising performance accuracy during both the training and testing phase of the PST. In other words, rTMS did not simply shift performance along a speed-accuracy trade-off [44], but instead improved both speed and precision to obtain reward, as measured by the subject’s reward rate (i.e., the number of correct responses produced per second of task activity). These findings dovetail neatly both with computational theories of tonic dopamine functioning and with pharmacological studies of dopamine and learning [45, 46].

In particular, theoretical and computational work have largely concentrated on phasic dopaminergic signaling in predictive learning [16, 47], yet dopamine neurons operate in both a phasic and a tonic mode [46]. Accordingly, a recent computational theory of tonic dopamine, the ‘tonic dopamine hypothesis’, holds that the net rate of reward (or reward rate) – the critical determinant of response rates across various reinforcement learning task – might be represented by tonic levels of dopamine in the striatum [48]. Notably, pharmacological manipulations that enhance striatal dopamine release (e.g., sulpiride, cabergoline) or reduce reuptake (e.g., methylphenidate) are commonly seen in the vigor of ongoing behavior, rather than in learning processes [41, 49]. For instance, Westbrook and colleagues (2020) demonstrated that reaction time decreased when participants were administered sulpiride – a D2 receptor antagonist – in an effort-based reinforcement learning task, and that the effect of sulpiride on response speeding was larger for those with lower caudate dopamine synthesis capacity as measured by PET[49]. To note, the caudate, and not putamen, was the basal-ganlia region that displayed an increase in dopamine following rTMS [3]. Further, in both animals and humans, dopaminergic stimulation increases the allocation of effortful force for reward without trading speed for accuracy [50–52]. By contrast, patients with Parkinson’s disease (depleted dopamine) show a reduced ability to increase movement speed in response to reward incentives [53, 54]. These accounts have inspired a new set of modeling investigations, in which the rate of reward acts as an opportunity cost for time, thereby penalizing sloth, and is suggested as being coded by the tonic (as distinct from the phasic) levels of striatal dopamine [45, 46]. For example, a recent computational model demonstrated that motivation by reward can break the speed-accuracy trade-off, apparently by simultaneously invigorating movement and improving response precision [54]. In particular, the authors found that applying a noise-reduction cost to optimal motor control predicted that reward motivation can increase both velocity and accuracy, and it has been suggested that reward might potentially exert its effects on vigor of response via dopamine[54].

Because the tonic level of dopamine is hypothesized to change very slowly over time, it is plausible that the continuous application of rTMS (15 minutes) may have had more of an accumulating effect on tonic levels of dopamine in the striatum. If tonic dopamine really does convey the net reward rate and enhance response vigor, as proposed by the ‘tonic dopamine hypothesis’, it is clear why higher levels of dopamine (as a result of 10-Hz rTMS administration) would result in overall faster responding in the PST and higher reward rate in the training phase of the task. Together, these behavioral and computational findings aim to capture the coupling between phasic and tonic dopamine, and help explain results linking vigor to predictions of reward. We hope these finding will motivate future investigations.

In sum, we demonstrated that stimulating the left DLPFC using a 10-Hz rTMS protocol, which has been shown to potentiate dopamine release in the dorsal striatum[3], is capable of enhancing the reinforcement learning function of the dorsal striatum, as captured by changes in reward rate (whose neural underpinning has been suggested to be tonic levels of dopamine in the striatum), and a computational bias towards learning from positive RPE signals (whose neural underpinning has been suggested to be phasic levels of dopamine in the striatum). Although the study did not directly measure dopamine release following 10-Hz repetitive TMS, the behavioral and computational findings are well positioned to provide indirect evidence of the effect of TMS on both a tonic and phasic mode of striatal-dopamine functioning. While more research is needed, combining rTMS with behavioral and computational measures of reinforcement learning opens new avenues for basic research on reinforcement learning and clinical studies of several psychiatric disorders that involve cognitive and behavioral disturbances attributed to disrupted dopamine signaling in the striatum.

## Acknowledgements

We thank Mei-Heng Lin, Malte Gueth, Galit Karpov and Marissa Cortright for their assistance with data collection.

## Author contributions

T.E.B. and K.B designed the research; K.B and S.C. collected the data; K.B. and T.E.B performed behavioral and computational analysis; C.M. and J.C. provided support in computational modelling; T.E.B. and K.B. wrote the first draft of the paper; all authors contributed to the manuscript.

## Funding

This work was supported by the National Institute on Drug Abuse of the National Institutes of Health [Award Number 1R21DA049574-01A1].

## Footnotes

a Ott and colleagues (2011) previously tested this hypothesis using prefrontal continuous theta burst stimulation (cTBS), a TMS protocol shown to decrease both motor cortex activity and free dopamine in the striatum [23]. The authors revealed no influence of cTBS on PST learning performance, but left-hemispherical DLPFC stimulation led to a more reward-guided performance during the testing phase of the task relative to a right-hemispherical cTBS group and Sham (vertex cTBS) group. fMRI data also revealed an increased model-derived RPE signal in the ventral striatum [21]. However, while these findings where initially attributed to a cTBS-induced increase in dopamine activity, a corrigendum [22] corrected this statement given that PET-TMS evidence indicated a decrease of free dopamine in the striatum following cTBS [23]. Further, the inhibitory effects of cTBS on neural activity [24] have recently been challenged [25, 26]. Thus, the effects of 10-Hz rTMS on reinforcement learning, as measured by the PST and Q-learning, remain unknown.

b The original PST includes a third difficult pairing (EF, rewarded 60% and 40% respectively), for which participant accuracy is often at or near chance; here, to reduce the duration of the task, and make it easier for subjects to learn, we removed the EF pair from the stimulus set.

## Notes

### Competing Interest Statement

The authors have declared no competing interest.

